# Regulation of neurogenesis and gliogenesis by the matricellular protein CCN2 in the mouse retina

**DOI:** 10.1101/2021.04.01.438112

**Authors:** Golam Mohiuddin, Genesis Lopez, Jose Sinon, M. Elizabeth Hartnett, Anastasiia Bulakhova, Brahim Chaqour

## Abstract

Cellular communication network (CCN) 2 is an extracellular matrix protein with cell type- and context-dependent functions. Using a combination of mouse genetics and omic approaches, we show that CCN2 is expressed in early embryonic retinal progenitor cells (RPCs) and becomes restricted to fully differentiated Müller glial cells (MGCs) thereafter. Germline deletion of CCN2 in mice decreases BrdU labeling, reduces RPC pool, and impairs the competency of remaining RPCs to generate early and late born retinal cell types. Retinal hypocellularity and microphthalmia ensue. The transcriptomic changes associated with CCN2 inactivation include reduced marker and transcriptional regulator genes of retinal ganglion cells, photoreceptors and MGCs. Yap (Yes-associated protein), a singular node for transcriptional regulation of growth and differentiation genes, is also a target of CCN2 signals. In an organotypic model of *ex vivo* cultured embryonic retinas, CCN2 and YAP immunoreactivity signals overlap. Lentivirus-mediated YAP expression in CCN2-deficient retinal explants increases the number of differentiating Sox9-positive MGCs. Taken together, our data indicate that CCN2 controls the proliferative and differentiation potentials of RPCs ultimately endowing, a subpopulation thereof, with Müller glial cell fate.

**Summary statement:** A CCN2-YAP regulatory axis controls retinal progenitor cell growth and lineage commitment to neuronal and glial cell fates.

## Introduction

The fully developed mammalian retina is composed of six principal types of neurons and the Müller glia cells (MGCs), all of which appear in a fixed but somewhat overlapping chronological order. The expression of specific transcription factors (TFs), or a combination thereof, plays a critical role in directing RPCs to adopt specific retinal cell fates (Vetter and Brown, 2001). However, little is known about the environmental extrinsic factors that control RPC fate decisions.

One set of extracellular signals originates from the retinal extracellular matrix (ECM), the acellular, protein-rich scaffold in which cells and tissues are embedded (Ishikawa et al., 2015; Reinhard et al., 2015). With the advent and success of cutting edge omic methodologies, the composition of the retinal “matrisome” became well known (Collin et al., 2019). Of the well characterized ECM proteins, Fibronectin, Collagen IV and Laminins, the main ECM components of basement membranes including Bruch’s membrane and the inner limiting membrane (INL), have been shown to play a critical role in optic cup morphogenesis (Kwan, 2014). Serjanov et al demonstrated that β2-containing laminins provide epigenetic cues that regulate retinal progenitor cell (RPC) cytokinesis and fate commitment (Serjanov et al., 2018). Conversely, deletion of the interphotoreceptor matrix (IPM) proteoglycan 1 (IMPG1) and/or IMPG2 caused no evident developmental or morphological changes in the retina, but led to the formation of subretinal lesions and reduction of visual function (Salido and Ramamurthy, 2020). However, the functional importance of many other ECM components in retinogenesis remains poorly understood. Our recent studies identified the matricellular protein cellular communication network 2 (CCN2) aka connective tissue growth factor (CTGF) as a critical component of the retinal matrisome essential for retinal angiogenesis (Lu et al., 2020). Herein, we demonstrate that CCN2 is also an important component of the RPC secretome and further determine its importance in retinogenesis.

CCN2 is a multimodular ECM protein expressed in a restricted pattern during development of the central nervous system (CNS) as well as in non-neural tissues (Heuer et al., 2003). During development, a strong neuronal expression was present especially in the microtubule-associated protein 2 mRNA-positive neurons in the cortical layer VII and the dorsal endopiriform, and in the deep layers of the olfactory glomeruli and accessory olfactory nucleus (Heuer et al., 2003). However, although CCN2 functions extend to developmental processes in the CNS, a mechanistic understanding of its role in neurogenesis is an important gap in the rigor of prior knowledge.

For a while, CCN2 association with tissue scarring and fibrotic diseases has provided the impetus for many investigations of CCN2 biology and biochemistry (Abraham, 2008; Brigstock, 2003; Chintala et al., 2012). The primary translational product of the CCN2 gene is a secreted ∼40 kDa protein organized into four distinct structural modules that exhibit homology to conserved regions in a variety of extracellular mosaic proteins including insulin-like growth factor binding protein domain, von Willebrand factor type C repeat, thrombospondin type 1 repeat and cysteine knot motif (Chaqour and Karrasch, 2020; Krupska et al., 2015). The intact protein functions in a variety of tissues and organs through interactions between its modular structures with integrins (e.g., α_v_β_3_ integrin), non-integrin receptors (e.g., p75^NTR^, TrkA), and growth factors (e.g., nerve growth factor, bone morphogenic protein-2, transforming growth factor (TGF)-β, neurotrophins) (Lau, 2016). However, CCN2 effects are cell type-specific and context dependent. For instance, CCN2 functions to promote survival of olfactory bulb neurons, but it induces apoptosis of immature neurons via TGF-β2–SMAD signaling (Khodosevich et al., 2013).

This study was designed to determine the expression, regulation, and function of CCN2 during retinogenesis. We report for the first time dynamic expression of the CCN2 gene in RPCs and glial cell precursors. We show that loss of CCN2 function produced hypomorphic conditions subsequent to alterations of transcriptional and posttranscriptional regulators and markers of growth and differentiation of various retinal cells types. The effects of CCN2 deletion can be rescued, at least in part, through ectopic expression of YAP, a transcriptional co-activator targeted by CCN2 signals. Our studies reveal novel ECM-derived developmental signals required for neurogenic and gliogenic switches during the temporal differentiation programs in the retina.

## Results

### Expression pattern of CCN2 in the developing and adult retina

We previously showed that CCN2 is expressed in the retinal vasculature as it emerges from the optic nerve head in newborn mice and plays a critical role in vessel homeostasis and barrier function in adult mouse retina (Moon et al., 2020). Since CCN2 exhibits a widespread expression in the CNS (Hertel et al., 2000; Krupska et al., 2015), we examined its potential expression in the avascular retina during early and late embryonic stages. Quantitative (q) PCR-based analysis of CCN2 expression showed high CCN2 mRNA levels in embryonic retinas harvested at E12, E14, and E16 although the signal appears to subside around the time of birth (Fig. 1A). To study in more details the expression of the CCN2 gene during retinal neurogenesis, we used CCN2-enhanced green fluorescent protein (GFP) transgenic mice wherein a 100-kb CCN2 promoter-GFP bacterial artificial chromosome transgene was inserted into the mouse genome (Gong et al., 2003). We further analyzed GFP expression as a proxy for endogenous CCN2 through direct visualization of GFP fluorescence in fixed retinal cryosections. A strong GFP fluorescence was detected at E11.5 in the outer neuroblastic layer (NBL) (Fig. 1B), which is largely dominated by RPCs. Fluorescence-activated cell sorting (FACS) of single cell suspension from enzymatically digested E14 retinas showed a population of GFP expressing cells (up to 4.5%) at E11.5 (Fig. 2C). To obtain a more detailed understanding of CCN2-GFP distribution, we examined the expression of the CCN2-GFP reporter at subsequent retinal embryonic stages. At E14, CCN2-GFP^+^ cells were still localized but scattered in the outer NBL (Fig. 1D and 1G). We noted that hyaloid vessels which are prominent in the embryonic eye were also CCN2-GFP-positive. As retinogenesis proceeded, (E16-E18), CCN2 expression became scattered in the expanding neuroblastic region (Fig. 1E-F and H-I). Further immunohistochemical analyses showed that from P1 onwards (i.e., P12), CCN2-GFP expression could not be detected in neurogenic cells and was largely restricted to isolectin B4-stained retinal vasculature and in Müller glia cell (MGC) body emplacement in the INL (Fig. 1J). Meanwhile, through mining of single cell RNA-Seq data of mouse embryonic retina reported by Lu et al (Lu et al., 2020), we found that CCN2 was expressed in several RPC clusters during retinogenesis (Fig. S1-A-B), which corroborates our findings. Sox2 expression, which universally marks neural stem and progenitor cells throughout the CNS including the neural retina (Taranova et al., 2006), was associated with 45% of CCN2-GFP^+^ cells at E14 (Fig. S2-C and S2-E). The expression of CHX 10, one of the earliest RPC markers (Clark et al., 2008), was associated with 23% of CCN2-GFP^+^ cells (Fig. S2-D-E). An overview of retinal sections at E14 showed that nearly 50% of CCN2-GFP^+^ cells were proliferating Ki67^+^ cells (Fig. S2F-G). Thus, CCN2 signals could be reiteratively used for cell proliferation and specification decisions during retinogenesis.

**Figure 1.**
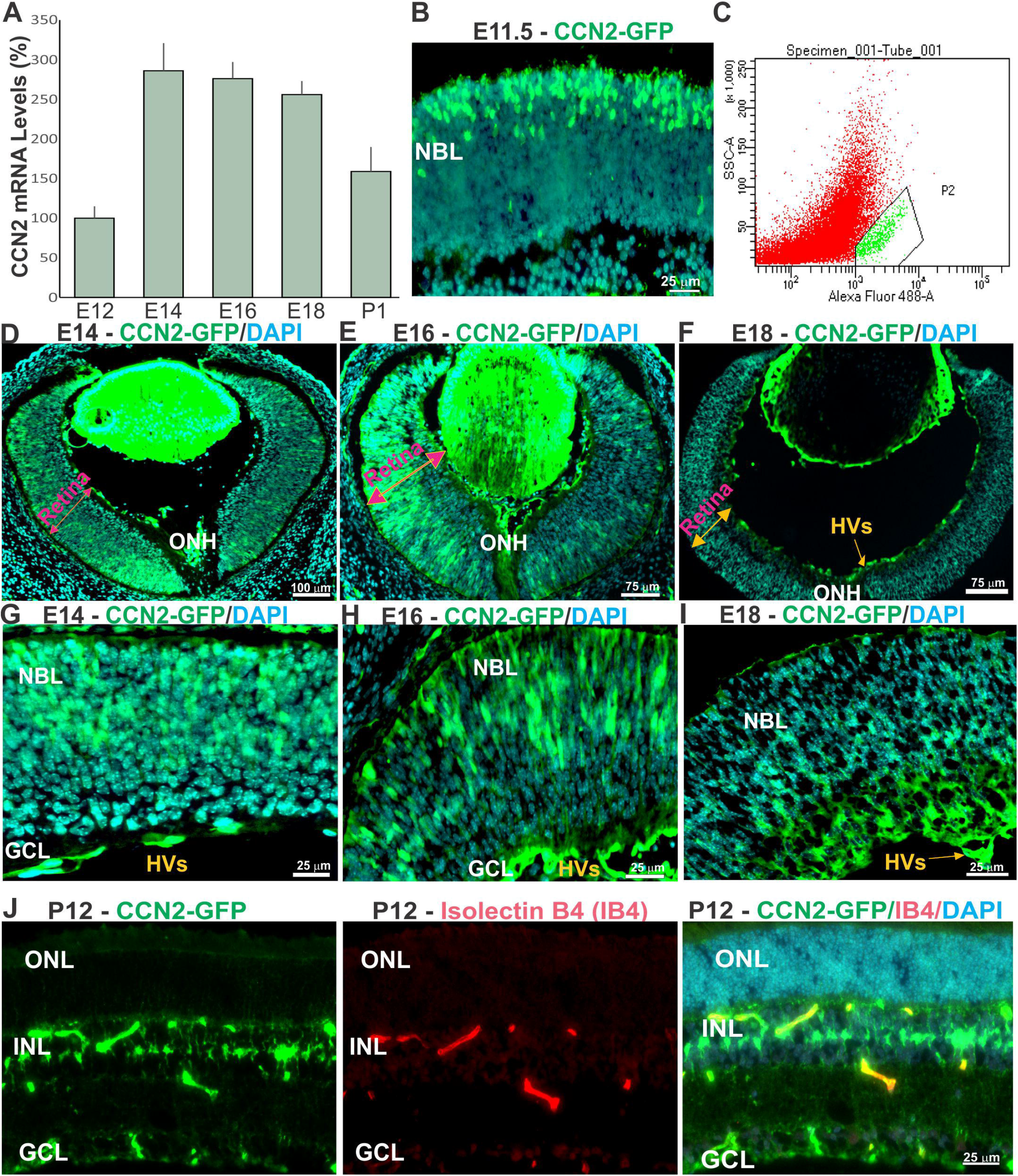
Expression pattern of CCN2 in embryonic mouse retina. (A) CCN2 transcript levels during early stages of retinogenesis. CCN2 mRNA levels were determined by qPCR and normalized to those of 18S rRNA (n=5). (B) Transverse histological section of E11.5 CCN2-GFP mouse retinas stained with DAPI for nuclear localization of retinal cells. Note that CCN2-GFP^+^ cells were localized in the outer neuroblastic layer (NBL). (C) Quantification of CCN2-GFP^+^ cells in transgenic mice expressing GFP under the CCN2 promoter control through FACS analysis dot plots representing the percentage population of RPC^+^ CCN2-GFP cells at E11.5. (D-I) Dynamic changes in CCN2-GFP expression and localization during embryonic eye formation in CCN2-GFP mice. CCN2-GFP expression was directly visualized by immunofluorescence microscopy of fixed cryosections, collected in the plane of sectioning through the optic nerve head (ONH). High magnification images of the same retinas are shown in G-I panels. HVs denote positive CCN2-GFP staining of the hyaloid vasculature. (J) Transverse retinal section of a P12 CCN2-GFP mouse retina showing colocalization of CCN2-GFP with either IB4-stained retinal vessels and INL cells characteristic of MGCs localization.

**Figure 2.**
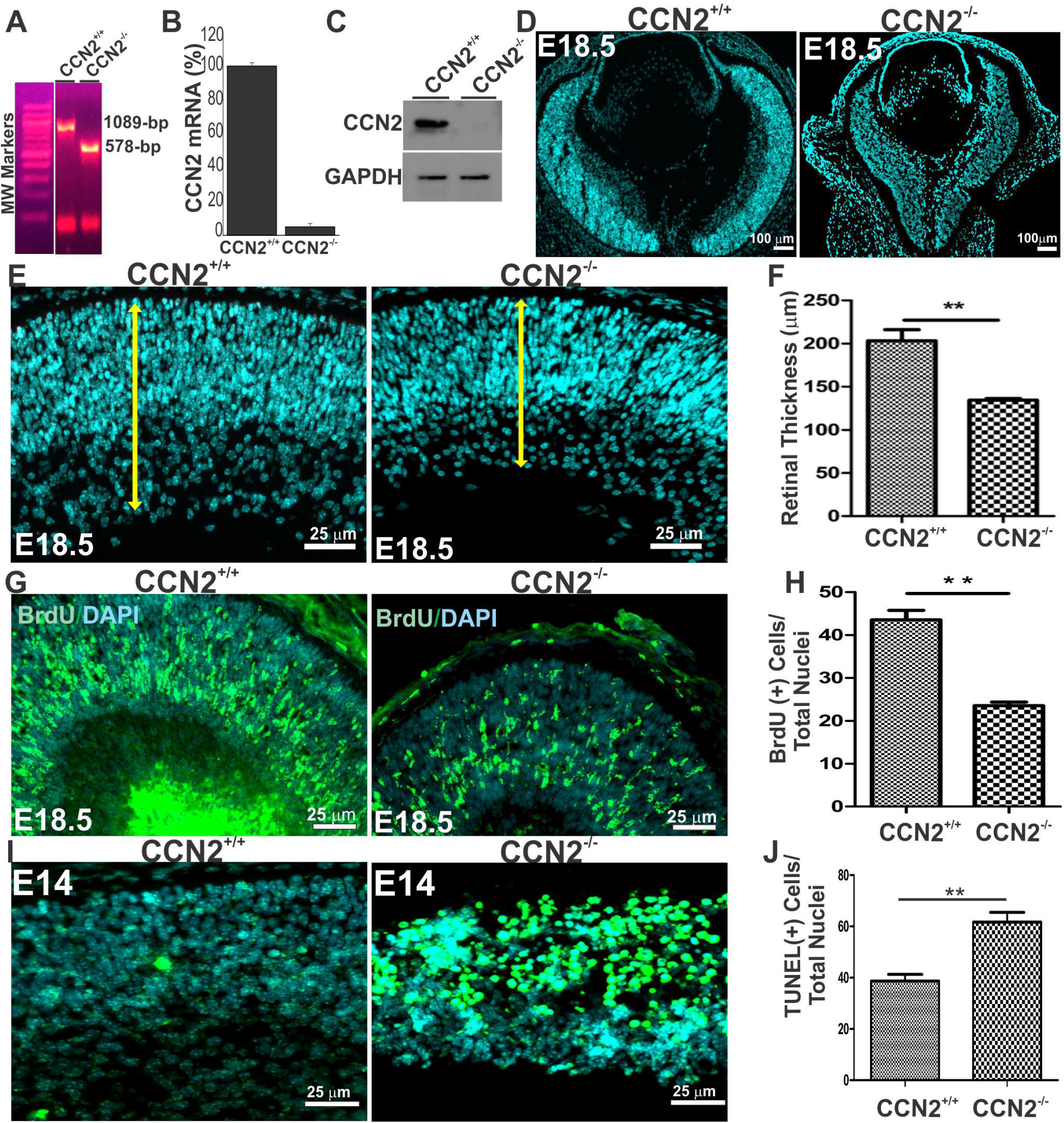
CCN2 deficiency resulted in retinal microphthalmia and hypocellularity defects. **(A-C)** Targeting the CCN2 genomic locus with the CCN2 flox-neomycin allele with CMV-Cre-mediated excision of loxP sites was detected by PCR as 1089- and 578-bp bands of WT mice and mutant CCN2^-/-^ alleles, respectively. (B) Relative CCN2 mRNA levels in lysates of WT and CCN2^-/-^ mouse retinas as determined by qPCR. Each measurement was performed in triplicate. Values are means ± S.E. *, p <0.001 *vs* CCN2^+/+^ (n=4). (C) Western blot analysis of protein lysates of WT and CCN2^-/-^ mouse retinas using CCN2 antibodies. GAPDH signal was used as a loading control. (D-F) Comparative morphometric analysis of transverse histological sections of DAPI stained WT and CCN2^-/-^ eyes at E18.5. Quantitative analysis of retinal thickness is shown in (F). Measurements were performed in the posterior retina near the optic nerve exit. (G-H) Effects of CCN2 deletion on cell proliferation during retinogenesis. Representative image of BrdU incorporation in transverse section of WT and CCN2^-/-^ mouse retinas. The average number of BrdU+ cells of three mouse retinas is shown in (H). Data are means ±S.E. **, *p* <0.01 *versus* WT. (I-J) Apoptotic cells were examined by TUNEL assay in retinal cross-sections of WT and CCN2^-/-^ mice. The average number of TUNEL^+^ cells is shown in (J). Data are means ±S.E. **, *p* <0.01 *versus* WT.

### CCN2 Deletion altered RPC proliferation and differentiation

As a secreted matricellular protein, CCN2 localizes both within the interstitial space and pericellularly due to its heparin-binding activity, which suggests that CCN2 exerts both autocrine and paracrine actions (Kireeva et al., 1997). To determine the significance of CCN2 expression during retinogenesis, we generated mice with global deletion of CCN2 by crossing CCN2 floxed and Cre driver allele mice. The Cre gene, which is under the transcriptional control of human cytomegalovirus (CMV) minimal promoter, is expressed before implantation during early embryogenesis (Schwenk et al., 1995). Consequently, Cre recombination results in deletion of the CCN2 gene in all cell types including stem and progenitor cells. CCN2 gene inactivation was confirmed by PCR screening of genomic DNA, and qPCR and western blot analyses of retinal lysates, all of which showed suppressed CCN2 gene expression at the genomic, mRNA and protein levels, respectively (Fig. 2A-C). As CCN2 constitutive deletion caused perinatal lethality (Ivkovic et al., 2003), wild-type (WT) (i.e., CCN2^+/+^) and CCN2-deficient (i.e., CCN2^-/-^) mouse embryos were harvested at E18.5 for analyses. CCN2^-/-^ mice displayed clear bilateral microphthalmia (small eyes) but no anophthalmia (absent eyes) as CCN2 deletion resulted in eyes 21.7± 5.4% smaller in size than CCN2^+/+^ eyes (Fig. 2D). Histological analyses of CCN2^-/-^ mice through nuclear staining visualization of coronal sections showed evidence of nuclear loss in mutant retinas (Fig. 2E) compared to WT retinas. The majority of CCN2-deficient mouse retinas were consistently thinner (−29.7± 7.5%) than WT retinas (Fig. 2F). In addition, loss of CCN2 function reduced the number of proliferating Bromodeoxyuridine / 5-bromo-2’-deoxyuridine (BrdU)^+^ cells by 46% (Fig. 2G-H) and increased the number of apoptotic cells by 20% (Fig. 2I-J) suggesting that CCN2 deficiency compromised cell proliferation and survival.

### Transcriptomic changes induced by CCN2 deletion during retinal neurogenesis

To gain insights into CCN2 mode of action on retinal neurogenesis, we hypothesized that CCN2 signals remodel global gene expression in the retinal neuroepithelium. To test this hypothesis, retinas were collected at E18.5 from WT and CCN2^-/-^ mice and used for genome-wide analysis by bulk RNA-sequencing (RNA-seq) of the murine retinal transcriptome. Single sequencing libraries were generated from pooled retinas of each mouse pup and three biological replicates of WT and CCN2^-/-^ mice were sequenced at an average of 8.7M paired-end reads (2 × 75 bp) resulting in 91.14% congruent paired mapped reads. The overall similarity among WT and knockout samples was assessed by the Euclidean distance between samples. As shown in Fig. 3A, WT samples showed shorter distances among samples of the same group (WT or CCN2^-/-^) indicating that samples of the same group are closely related to each other. Similarly, unsupervised hierarchical clustering showed a consistent clustering and clear segregation in the normalized gene expression profiles of WT and CCN2 mutant retina samples (Pearson’s correlation coefficient r = 0.94) (Fig. 3B). We identified 482 significantly changed genes (283 downregulated and 199 upregulated genes) in CCN2^-/-^ compared with CCN2^+/+^ retinas (absolute log2 fold change >0.5, p < 0.05) (Fig. 3C-D). Genes regulated by CCN2 encode proteins with a wide range of biological activities, likely reflecting the large diversity of cell types and circuits affected by CCN2 deletion in the retina. Cell proliferation, hypoxia, DNA repair, glycolysis, and inflammatory response genes were all differentially expressed between WT and CCN2-deficient mouse retinas (Fig. 3E and Fig. S2). CCN2 deficiency resulted in downregulation of neuronal cell differentiation (e.g., basic helix-loop-helix family member e22 (BHLHE22), neuropeptide-like 4, dopachrome tautomerase), growth factors (e.g., TGF-α, PDGF-B, BMP4), ion channels (e.g., CLIC6), basement membrane protein (e.g., COL8A1, LAMB1), cell-cell communication (e.g., delta like non-canonical Notch ligand 2 or DKK2), cellular stress response (e.g., serum/glucocorticoid regulated kinase 1 (or SGK1), transcription factors (e.g., JUN B, zinc finger protein 365, Azin2), numerous small and long non coding RNAs (e.g., SNORD58B, miR654, 2310010J17Rik) and cytoskeletal proteins (e.g., TTLL13).

**Figure 3.**
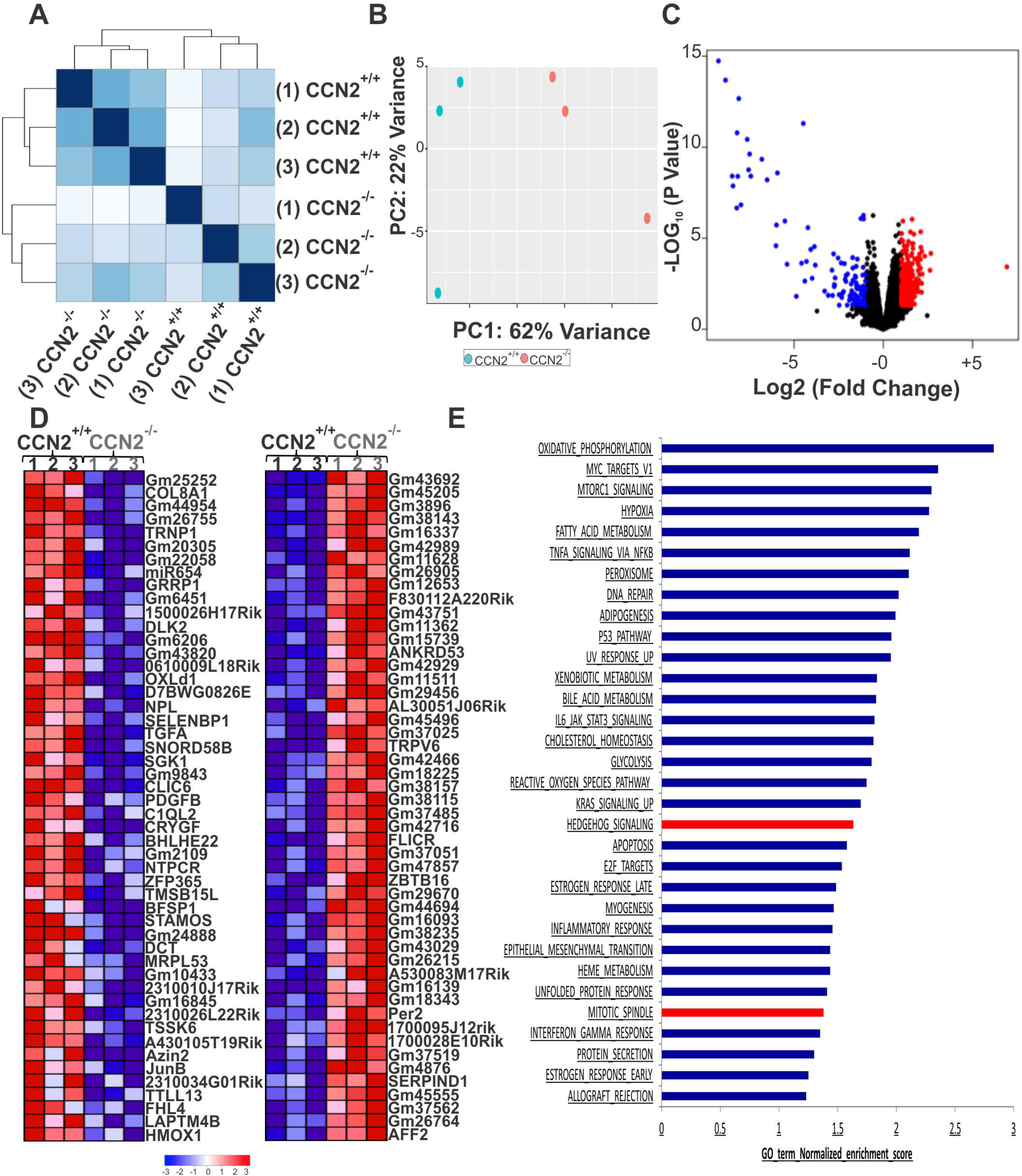
Deep transcriptome profiling of WT and CCN2^-/-^ embryonic mouse retinas. (A) Dendrograms of dissimilarity measures based on Euclidean distances using raw count data from three replicates of WT and CCN2^-/-^ mouse retinas. High correlations and low to medium variability among samples of the same group when compared to each other. (B) a Principal component analysis (PCA) of distinct hallmark gene expression profiles with clear segregation of WT from CCN2^-/-^ retinal transcriptomes of three distinct biological replicates, plotted in a two-dimensional volumetric space. (C) Volcano plot (significance *vs* fold change of significantly down-regulated (blue) and up-regulated (red) genes (fold change > 1.5 and P < 0.05) between E18.5 WT and CCN2^-/-^ retinas as determined by RNA-seq analysis. (D) Heat map and hierarchical clustering of 50 genes showing statistically significant changes in gene expression between WT and CCN2^-/-^ mice. Red indicates high expression and blue indicates low expression. (E) Pathway enrichment analysis of differentially expressed genes in CCN2^-/-^ *versus* WT mouse retinas. The clusters with functional terms that reached significance of p < 0.05 are shown. Online functional databases were used to extract each term.

We performed further data mining for the relevant function of the differentially expressed genes using Gene Set Enrichment Analysis (GSEA) (Subramanian et al., 2005) allowing all detected unfiltered genes to be mapped to defined gene sets (e.g., pathways), irrespective of their individual change in expression. We found that differentially expressed gene sets between the CCN2^-/-^ and CCN2^+/+^ groups mapped to Notch, Wnt and Hedgehog signaling pathways (Fig. 4A-D). Of these, Dll1, Dll3, Hes 1, Hes 5, Hey1 and Hey 1 of the Notch signaling pathway were particularly downregulated in the retina of CCN2-deficient mice. Components of the Wnt pathway such as LRP5, LRP6, Wnt2b were upregulated whereas Wnt ligands such as Wnt7a, Wnt7b, Wnt5a and Wnt9a were downregulated in CCN2^-/-^ compared to CCN2^+/+^ mouse retinas. Ptch and sufu which regulate sonic hedgehog (Shh) signaling were upregulated in CCN2^-/-^ *versus* CCN2^+/+^ retinas (Fig. 4C). These unalike transcriptomic alterations reflect the disparity of CCN2 effects on the different cell types in the retina. Interestingly, CCN2 deficiency significantly downregulated the expression of core component of the Hippo signaling pathway including YAP (Fig. 4D), commonly viewed as the main avenue where various pathways (e.g., Notch and Wnt) converge to control cell proliferation and differentiation in the embryo (Hansen et al., 2015). YAP immunoreactivity was found in the outer neuroblastic nuclear layer of WT mouse retina whereas CCN2-deficient retina showed little or no YAP immunoreactivity signals (Fig. 4E). Quantitative PCR analysis further validated differential expression of several components of the Hippo YAP pathway including YAP, Yes, LATS1, LATS2, NLK/Nemo-like kinase, AXN2, and AMOTL2 (Fig. 4F). CCN2 deficiency significantly reduced the transcript levels of NLK, which when depleted in a tissue, increased YAP phosphorylation and reduced YAP transcriptional activity (Moon et al., 2017). Thus, CCN2 deficiency recapitulates, at in part, a loss of YAP function phenotype.

**Figure 4.**
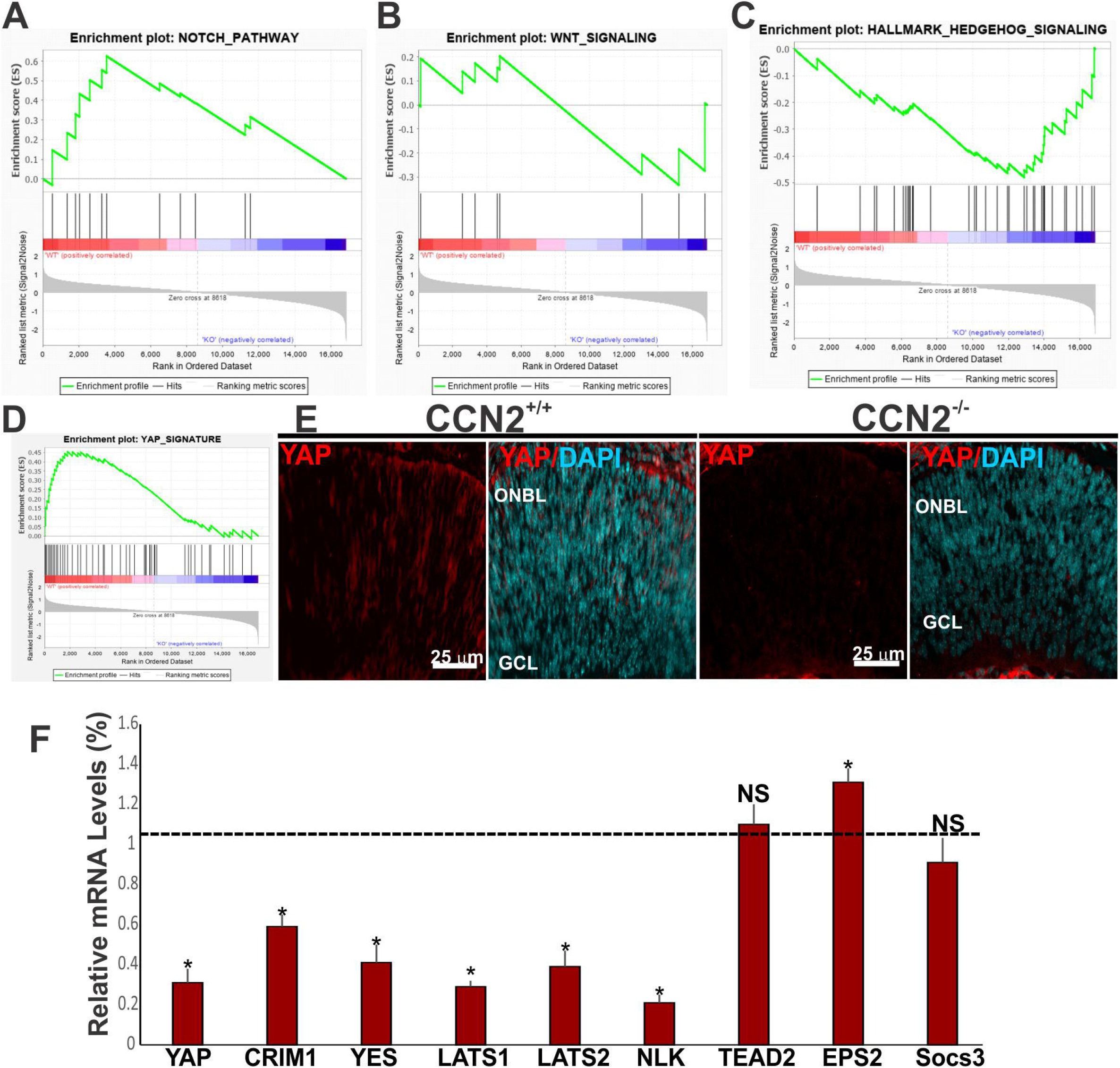
CCN2-dependent transcriptional program induced combinatorial signaling perturbations that include YAP Network Genes. (A-D) Pathway enrichment analysis of differentially expressed genes in CCN2^-/-^ versus WT mouse retinas with the gene set of Notch (A), Wnt (B), Sonic Hedgehog (C) and Hippo YAP signaling pathways. Online functional databases were used to extract each term. Genes whose expression levels are most closely associated with the CCN2^+/+^ group are located at the left, whereas genes from the CCN2^-/-^ gene set within the ranked list are located at the right. (E) Representative image of YAP immunostaining in transverse section of retinas of CCN2^+/+^ and CCN2^-/-^ mice. (F) Relative mRNA levels of YAP target genes in retinal lysates of CCN2^+/+^ and CCN2^-/-^ mice. The mRNA levels in CCN2^+/+^ mice were set to 1. Each measurement was performed in triplicate. (*p < 0.05, n = 3).

### CCN2 expression is crucial for the onset and propagation of retinal cell growth and differentiation

GSEA analysis showed that the transcriptomic changes associated with CCN2 deletion included a subnetwork of genes and pathways with the highest significance score that prioritized decreased growth and increased apoptosis (Fig. 5A-B). To identify the retinal cell types affected by these changes, we used qPCR to validate differential expression of selected gene markers including Sox2, which is largely responsible for neurogenic competence (Gorsuch et al., 2017), Pax6 which controls RPC multipotency by regulating cell type specification (Tao et al., 2020), and Chx10, which is expressed in proliferating retinal progenitor cells throughout retinal development (Green et al., 2003). As shown in Fig. 5C, Sox2 and Pax6 mRNA levels were significantly reduced in CCN2^-/-^ mouse retina further supporting our RNA-seq data. Similarly, Sox2 and Pax6 immunoreactivity in retinal transverse sections was markedly diminished in CCN2-deficient retinas compared to their WT counterparts (Fig. 5-D-E). CCN2 deficiency induced a more dramatic reduction of Sox2 (−30%) than Pax6 (−17%) mRNA levels indicating changes in the proper Sox2 to Pax6 ratio, which is key to the maintenance of RPC neurogenic competence and multipotency (Tao et al., 2020). Meanwhile, the expression of Chx10 was significantly reduced in CCN2^-/-^ mouse retinas as well (Fig. 5C and 5F), which is consistent with decreased proliferating RPC number in the mutant mice. Conversely, the mRNA levels of Six6 (aka Optx2) homeodomain factor, a strong tissue-specific repressor, that compromises RPC survival (Toy et al., 2002) was increased in CCN2^-/-^ *versus* CCN2^+/+^ retinas. Increased Six6 expression is consistent with the role of Six6 as a repressor cyclin-dependent kinase inhibitors (Li et al., 2002).

**Figure 5.**
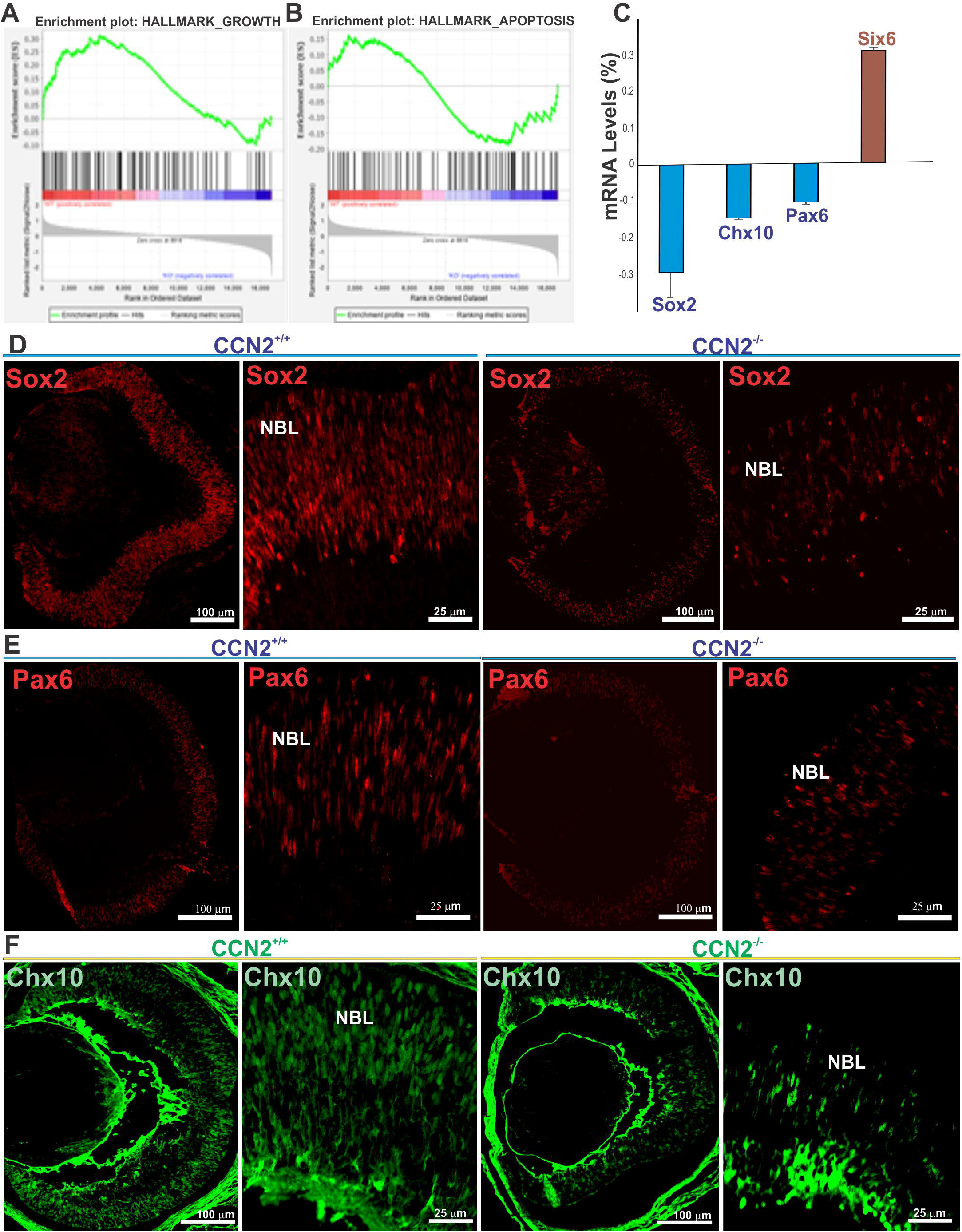
CCN2 deficiency alters RPC growth, survival and differentiation. (A-B) GSEA of mouse retinas showing upregulation of growth and apoptosis signature genes in WT compared to CCN2^-/-^. The graphs display the enrichment gene set based on pathways provided by the hallmark Msig database showing upregulation of growth signature genes in CCN2^+/+^ compared to CCN2^-/-^ mouse retinas (A) while a majority of apoptosis genes were upregulated in CCN2^-/-^ compared to CCN2^+/+^ retinas (B). (C) Relative mRNA levels of RPC markers in retinal lysates of CCN2^+/+^ and CCN2^-/-^ mice. The mRNA levels in CCN2^+/+^ mice were set to 1. Each measurement was performed in triplicate. (*p < 0.001, n = 3). (D-F) Representative image of Sox2, Pax 6 and Six6 immunostaining in transverse section of retinas of CCN2^+/+^ and CCN2^-/-^ mice.

### CCN2 deletion alters RPC fate specification in the retina

Differentiation of six neuronal and one glial cell types from RPCs follows a loose temporal sequence wherein retinal RGCs are differentiated first, followed by overlapping phases of development for cone photoreceptors (PRs), horizontal cells, amacrine cells, rods, and bipolar and MGCs. The differentially expressed gene sets between WT and CCN2^-/-^ retinas identified by GSEA analysis included RGC, PR- and MGC-specific genes which showed concordant differences between and CCN2^+/+^ and CCN2^-/-^ mouse retinas. We used qPCR to validate the expression of selected retinal cell type-specific genes. For RGCs, the expression of Brn3/Pou4f1 known to promote RPC-differentiation into RGCs and inhibit non-RGC differentiation gene programs (Moshiri et al., 2008), was significantly reduced in CCN2^-/-^ mouse retina (Fig. 6A). Brn3b/Pou4f2 immunostaining in retinal transverse sections was clearly detected in the migrating RGCs in the neuroblastic layer and the forming ganglion cell layer at the inner surface of the developing retina of WT mice (Fig. 6B). However, their number was significantly reduced in CCN2^-/-^ compared to CCN2^+/+^ retina (Fig. 6C). Similarly, the transcript levels of Foxp2, and Opsin 4 (Opn4) which are specifically expressed in a subset of RGCs (Mao et al., 2014), RNA-binding protein with multiple splicing (Rbpms), which is expressed in the entire RGC population (Kwong et al., 2010), g-synculin (Sncg), and neuritin 1 (NRN1) which are critical for RGC survival (Chintalapudi et al., 2016), were all downregulated as a result of CCN2 deletion. Conversely, the expression of Satb1 which was shown to be dispensable for initial RGC differentiation, and Satb2, which counteracts the effects of Satb1(Sweeney et al., 2019), was increased in mutant mouse retinas. The expression of Scn8a, which promotes RGC death (Alrashdi et al., 2019) was also upregulated in CCN2 mutant mouse retinas. We further analyzed the expression pattern of Islet-1&2, two important transcription factors important for retinal development. Islet-2 is considered as an early marker for a subpopulation of RGCs while Islet-1 labels amacrine, bipolar, and RGCs. As shown in Fig. 6D, Islet-1 &2-labeled cells were observed in the GCL of either CCN2^+/+^ or CCN2^-/-^ retinas although a subset of Islet-1&2-positive cells localized in the inner part of the INL where putative Islet1-positive amacrine cells are. These can be distinguished from Islet1-positive bipolar cells located in the middle to the outer portion of the INL. The number of Islet-1&2-positive cells was significantly reduced in CCN2-deficient retina compared to WT (Fig. 6E). Conversely, the number of cells in the putative INL (including amacrine and bipolar cells) was significantly increased in CCN2^-/-^ retinas compared to CCN2^+/+^ retinas (Fig. 6F). Quantitative PCR analysis of amacrine and bipolar cell-specific markers showed increased Islet1, whereas Islet 2 mRNA levels were decreased in CCN2^-/-^ retinas compared to CCN2^+/+^ retinas (Fig. S3-A). These changes are corroborated by increased expression of several makers of amacrine and bipolar cell subpopulations in CCN2^-/-^ retinas compared to CCN2^+/+^ retinas (Fig. S3-A-B). Taken together, these data support the idea that CCN2 deletion promoted a transcriptional program facilitating amacrine and bipolar cell differentiation and survival at the expenses of the RGC lineage.

**Figure 6.**
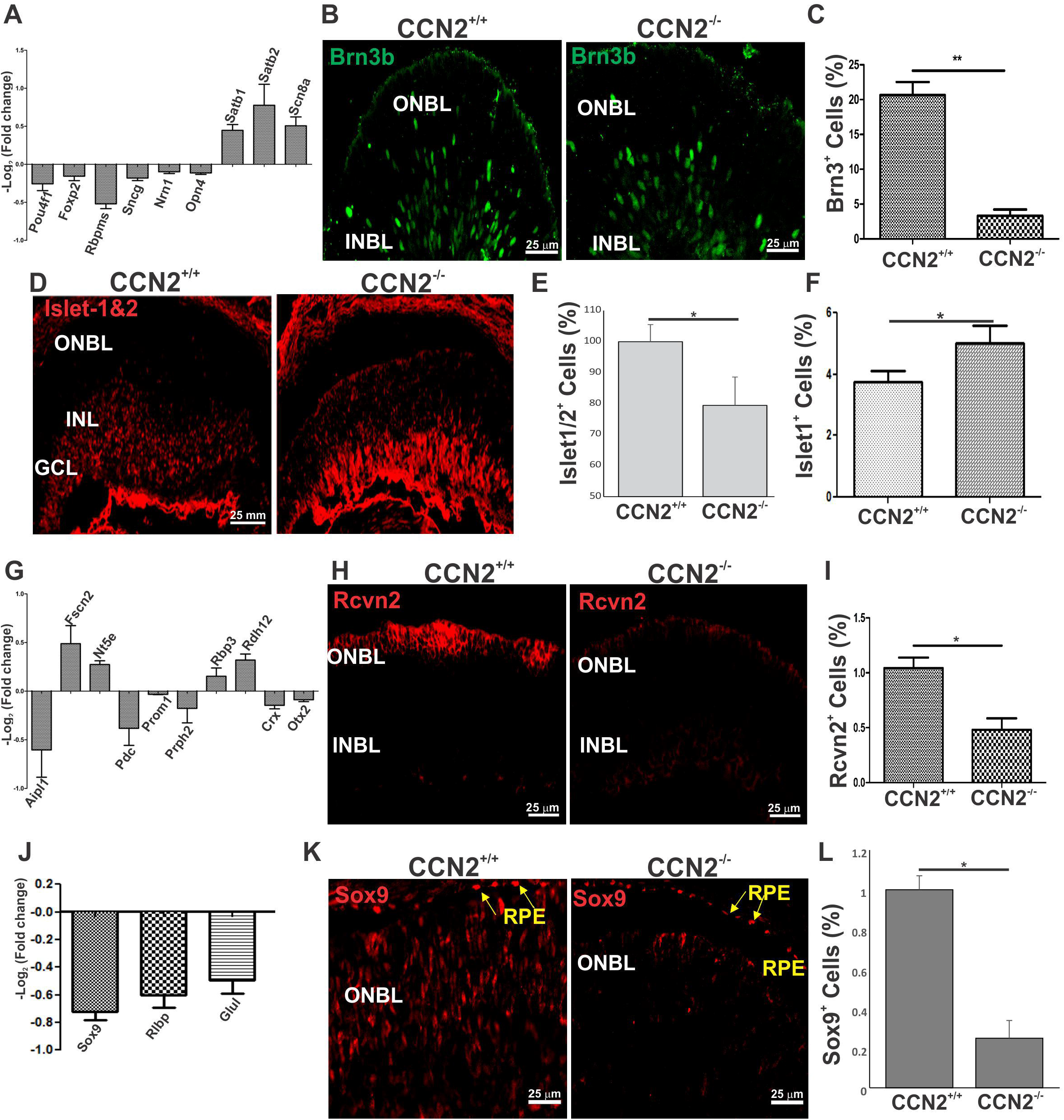
Effects of CCN2 deletion on molecular and phenotypical markers for RGCs, PRs, and MGCs. (A) (G) and (J) Relative mRNA levels of differentially expressed molecular markers for the RGC, PR and MGC phenotype in CCN2^-/-^ *versus* CCN2^+/+^ mouse retinas as determined by qPCR analysis. Each histogram represents the mean ± S.E. for triplicate samples. *, p< 0.05. (B), (D), (H) and (K) Representative image of Brn3b, Islet-1&2, Rcvn2 and Sox9 immunostaining in transverse sections of retinas of CCN2^+/+^ and CCN2^-/-^ mice. (C), (E), (F), (I) and (L) Quantitative analyses of Brn3b, Islet-1&2, Islet-1, Rcvn2 and Sox9 signals in transverse sections of CCN2^+/+^ and CCN2^-/-^ mice. *, p< 0.05.

As predicted by the RNA seq results, a reduced immunostaining with antibodies to recoverin2, a terminal differentiation marker of PRs was observed in CCN2-deficient mouse retinas (Fig. 6G-H). The number of recoverin2 immunoreactive PRs, which were confined to a few horizonal rows in the distal-most part of the neuroblastic layer of the CCN2^+/+^ mouse retina, was significantly reduced in CCN2^-/-^ mouse retina (Fig. 6I). Quantitative PCR analysis also confirmed a significant downregulation of Crx and Otx2, which are critical for RPC fate commitment and differentiation into cones and rods (Fig. 6G). Similarly, cone PR-encoding genes critical for the formation of the outer segment such as peripherin 2 (Prph2/Rds), and phosducin (Pdc) were downregulated as a result of CCN2 deletion. Non-retina specific transcript molecules such as fascin 2 (Fscn2), an actin-binding protein, and Nt5e, a cell surface antigen, were upregulated in the CCN2^-/-^ mouse retina. Thus, although CCN2 deletion resulted in depletion of a large PR cell population, small subpopulations of cone PRs may be increased in CCN2^-/-^ retina.

The decline in expression of RPC markers like Sox2 in CCN2-deficient retina is predictive of a subsequent alteration of MGCs, a non-neuronal cell type in which Sox2 expression is maintained (Gorsuch et al., 2017). Both MGCs and RPCs co-express many transcription factors, neurofilament proteins, and signaling pathway molecules. Of these, Sox9, GLUL, and RLBP1, which have been dubbed as MGC-specific molecular markers (de Melo et al., 2016), were significantly downregulated in CCN2^-/-^ *versus* CCN2^+/+^ retinas (Fig. 6J). Although Sox9 is also present in adjacent RPE cells, Sox9-positive cells in the ONBL were easily identifiable by their location (Fig. 6K). However, fewer Sox9-positive cells were detected in the NBL of CCN2^-/-^ mice and their number was significantly reduced in CCN2 mutant mouse retinas compared to those of WT mice (Fig. 6L).

### Gain of CCN2 or YAP function increased RPC differentiation into PRs and MGCs

Our transcriptomic studies identified YAP, a key singular node of progenitor cell growth and differentiation, as one of the important targets of CCN2 signals during neurogenesis. To delineate the role of YAP in CCN2-dependent regulation of retinogenesis, we used an organotypic model of *ex vivo* cultured embryonic retinas. Cultured retinal explants are known to develop very similarly to a retina *in vivo* and generate several retinal cell types that will migrate to the appropriate layer (Cayouette et al., 2001). We first analyzed the expression and distribution of CCN2 and YAP in this model using retinas explanted from the CCN2-GFP reporter mice. Retinal explants were prepared from embryos at E14 and cultured with PR facing down in a 6-well culture plate. Immunohistochemical analysis showed that after 1 week of culture, the CCN2-GFP signal largely overlapped with YAP immunoreactivity in the neuroblastic layer (Fig. 7A). Of the CCN2-GFP expressing cells, 71% were YAP-positive as well (Fig. 7B) confirming a potential regulatory relationship between CCN2 and YAP gene expression. Therefore, we performed a CCN2 gain of function and YAP rescue analyses through lentivirus (Lnv)-mediated expression of either the CCN2 or YAP cDNA into CCN2^+/+^ and CCN2^-/-^ retinal explant cultures. A lentiviral vector expressing the luciferase gene was used as a control transduction vector. Thereafter, transverse retinal frozen sections were stained for different markers and positively labeled cells were quantified. As shown in Fig. 7C-D, a small number of Rcvn2-immunoreactive cells could be seen in the ONBL of Lnv-luc-transduced CCN2^+/+^ and CCN2^-/-^ retinal explants. Conversely, immunoreactivity to Rcvn2 was widespread in both Lnv-CCN2-transduced CCN2^+/+^ and CCN2^-/-^ retinal explants wherein a significantly increased number of Rcvn2 immunoreactive cells could be identified in all depths of the neuroblastic layer (Fig. 7D). Similarly, the number of Rcvn2-postive cells was increased in ONBL of Lnv-YAP-transduced CCN2^+/+^ and CCN2^-/-^ mouse retinas (Fig. 7E-F) suggesting that CCN2 and YAP are, at least in part, redundant in cone PR differentiation. We next examined the retinal subtypes of CCN2-GFP expressing cells by immunostaining for Sox9, whose expression becomes downregulated in differentiating neuronal populations, but persists in MGCs into adulthood. As shown in Fig. 7G, a near overlap between CCN2-GFP and Sox9-positive cells was observed at E18.5 retinas from CCN2-GFP mice wherein ∼90% of CCN2-GFP-positive cells were Sox9-positive cells as well (Fig. 7H). Concordantly, ectopic expression of either CCN2 or YAP significantly increased the number of differentiating Sox9-positive MGCs whose total number increased by 80 and 70% in Lnv-CCN2 and Lnv-YAP transduced mouse retinas (Fig. I-J), respectively. Thus, CCN2-mediated YAP expression is essential for the differentiation and/or survival of differentiating MGCs.

**Fig. 7.**
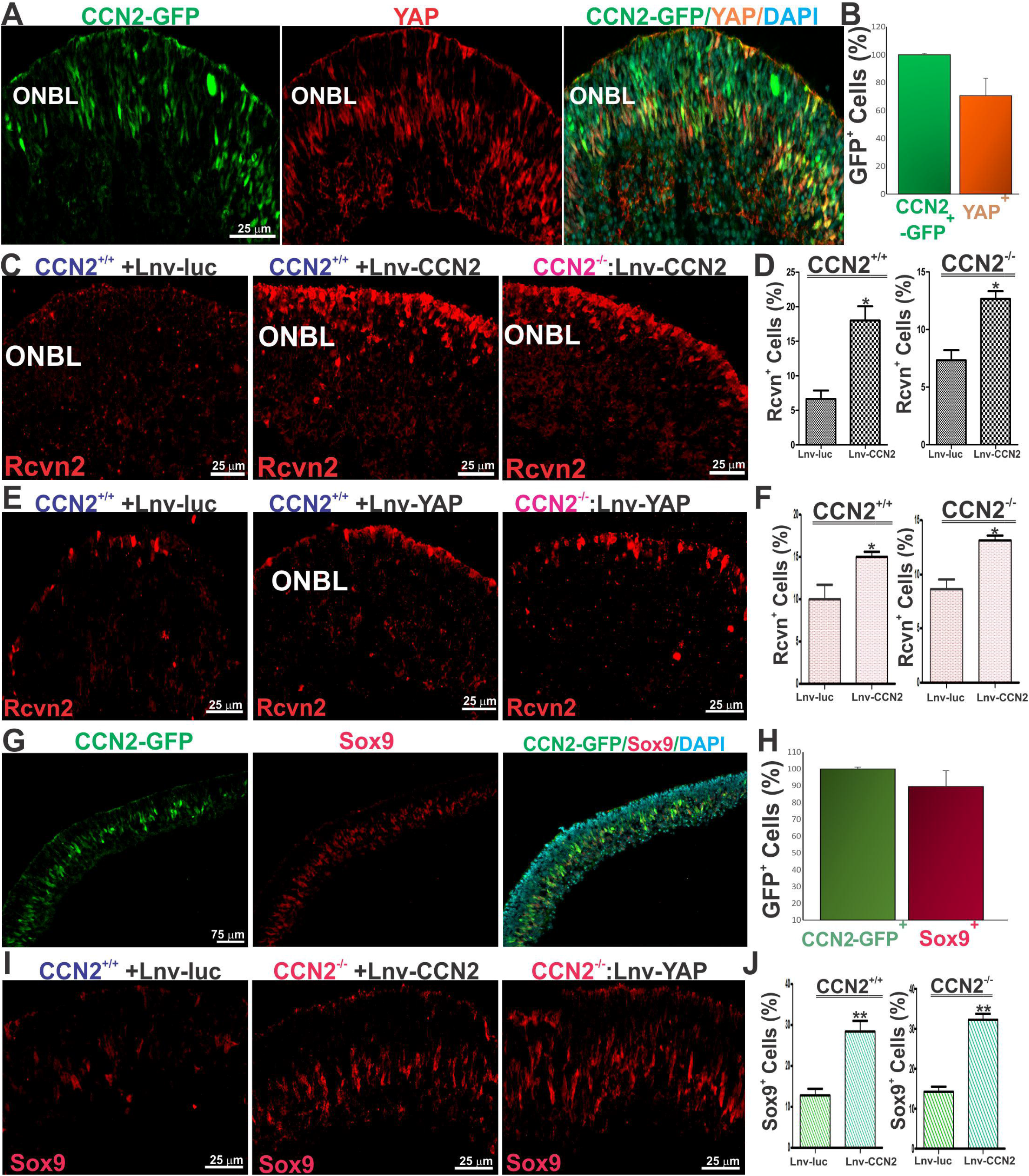
Gain of CCN2 or YAP function increased RPC differentiation into PRs and MGCs in ex vivo retinal culture explants. (A-B) Colocalization of CCN2-GFP and YAP signals in a CCN2-GFP mouse explanted at E14 and cultured for 6 days in a culture dish. The percentage of CCN2-GFP and YAP double positive cells is shown in (B). (C-F) CCN2^+/+^ and CCN2^-/-^ retinal explants were transduced with lentiviral vectors encoding either luciferase control (Lnv-luc) or the CCN2 or YAP transgene and further cultured for 6 days. Frozen sections were made and stained for Rcvn2 (shown in (C) and (E). The percentage of Rcvn2-positive cells are shown in (D) and (F). *, p<0.01. (G-H) Colocalization of CCN2-GFP and Sox9 signals in a CCN2-GFP mouse retinas explanted at E14 and cultured for 6 days in a culture dish. The percentage of CCN2-GFP and Sox9 double positive cells is shown in (H). (I-J) Effects of CCN2 and YAP on Sox9-positive MGC differentiation in CCN2^+/+^ and CCN2^-/-^ retinal explants. The percentage of CCN2-GFP and Sox9 double positive cells is shown in (J). *, p<0.01.

## Discussion

CCN2 is a multifunctional matricellular protein composed of several modules containing binding sites for integrins, low-density lipoprotein receptors, transmembrane proteoglycans, growth factors, ECM proteins and proteases (Lau, 2016). Through these interactions, CCN2 modulates various signaling pathways and induces subsequent changes in gene expression programs that culminate into activation of cellular processes such as cell adhesion, differentiation, proliferation and ECM protein remodeling. Despite the multimodular homology between CCN2 and other CCN proteins (Chintala et al., 2015), CCN2 exhibits unique cell type- and context-specific functions dependent on the availability of various receptors and binding partners in the cell membrane and extracellular environment. Since most CCN2 interacting partners (e.g., integrins, TGFβ-1, BMP, VEGF and Wnt) are implicated in CNS development, we could predict that CCN2 mediates multiple cellular events essential for retinogenesis.

First, our data showed that at the early stages of retinal neurogenesis, a subset of RPCs strongly expressed CCN2 which, subsequently became relocalized to the INL and no cells of the outer neuroblastic layer maintained CCN2 expression by the end of the gestational period. This spacio-temporal expression pattern might indicate that the CCN2 gene is expressed in the same cells and during their subsequent early differentiation phases or it might be expressed in distinct populations of retinal cells before they exhibit their cell fate biases. Of note, one third of CCN2 expressing cells co-expressed the RPC marker, Sox2, which regulates the choice between maintenance of RPC identity and differentiation (Taranova et al., 2006). As previous studies showed that exposure of embryonic mouse neural precursor cell cultures to recombinant CCN2 protein increased the number of Sox2-positive cells (Mendes et al., 2015), CCN2 may act as an upstream regulator of RPC-specific differentiation genes. Similarly, a proportion of CCN2 expressing RPCs was also positive for Chx10, which is largely expressed in early progenitors throughout retinal development. Chx10 has at least two essential roles: one in promoting RPC proliferation, and a second role in inducing bipolar cell differentiation (Green et al., 2003). Accordingly, nearly 50% of CCN2 expressing cells were proliferating suggesting a potential role of CCN2 in RPC proliferation as well.

Concordantly, CCN2 deletion induced gross retinal anatomical defects as mutant CCN2-deficient mice exhibited microphthalmia. This phenotype is reminiscent of that linked to defective levels of transcription factors essential for intrinsic control of mammalian retinogenesis such as Chx10 or Sox2. Previous studies have shown that even hypomorphic levels of Sox2 caused diverse microphthalmic phenotypes in postnatal animals (Karl et al., 2008). As in Sox2-deficient mice, CCN2 deletion led to a reduction in total cell counts due to reduced cell proliferation and increased cell apoptosis. Normal levels of CCN2 in WT animals were associated with RPC developmental potency and the ability to proliferate and differentiate, while CCN2 deletion resulted in their reduced proliferative capacity and differentiation culminating into retinal hypocellularity and microphthalmia.

Germinal deficiency of CCN2 resulted in major transcriptomic changes in the retina including alteration of gene expression programs for transcription factors involved in RPC specification and differentiation (e.g., Pou4f1, Foxp2, Rbpms, Sncg, Nrn1 and Opn4). Brn3b/ Pou4f1, which has been reported to be essential for RGC formation in mice (Moshiri et al., 2008), was significantly reduced in CCN2-deficient retinas compared with WT mouse retinas. Our GSEA analysis further showed that CCN2 reduced RGC number potentially through upregulation of the Scn8a gene. Scn8a encodes the Nav1.6 voltage-gated sodium (Nav) channel isoform, which commonly colocalizes with the Na+/Ca2+ exchanger (NCX) and β-APP, an indicator of imminent neuronal degeneration (Mojumder et al., 2007). Nav1.6 expression was suggested to reverse the function of NCX, and result in an increased influx of damaging Na^+^ and Ca^2+^ ions leading to cell death (Craner et al., 2004). Thus, loss of CCN2 function may not only reduce the ability of RPCs to differentiate into RGCs, but also increase death of differentiated cells.

Moreover, CCN2 deletion also downregulated the expression of Crx and Otx2, which together with Nrl, are essential for the postmitotic differentiation, maturation, maintenance and function of PRs (Yamamoto et al., 2020). The expression of other PR differentiation markers such as Nr2e3 and Rxrg, was similarly downregulated upon CCN2 deletion; however, Prdm1/Blimp1, which controls the balance between PRs and bipolar cells by suppressing bipolar identity in most of the cells (Xiang, 2013), was upregulated. Along these lines, CCN2-deficient retinas expressed substantially lower levels of other cone precursor-enriched non transcription regulators such as peripherin 2, phosducin, <u>aryl hydrocarbon receptor interacting protein like 1</u>, and prominin 1. Yet, expression of other neurogenic factors typically associated with non-PR cell types including fascin 2, Nt5e, Rbp3 and Rdh12 was maintained. Taken together, our data indicated that CCN2 expression is essential for adequate expression of postmitotically acting transcription factors directing terminal differentiation, and maturation of the majority of PRs, although a subpopulation of this cell type may persist in CCN2^-/-^ retina.

Another important result of our study design relates to the expression of Sox9, which functions to initiate retinogenesis and continues to be expressed in differentiated MGCs (Kang et al., 2012). Our data showed that the expression of CCN2 and that Sox9 nearly completely overlapped at the prenatal stages of retinogenesis and Sox9 expression was dramatically diminished upon CCN2 deletion. Congruently, the number of Sox9^+^ cells was 70% lower in CCN2-deficient retina compared to the WT and this is consistent with a major function of CCN2 in MGC fate specification and maintenance. Sox9 expression commonly is considered to indicate a switch of neuronal progenitors from neurogenesis to gliogenesis. Indeed, a study by Poche et al (Poche et al., 2008) showed that Sox9 conditional ablation or knockdown led to loss of Müller cell marker expression. Reversibly, conditional Sox9 expression in mouse progenitor lineage where it normally was not expressed, increased their proliferation and induced macroglia marker expression, indicative of premature gliogenesis (Guven et al., 2020). Similarly, CCN2 deletion altered the expression of early glia markers such as Rlbp and Glul, which exhibited an expression pattern identical to that of Sox9 in WT *versus* CCN2-deficient mouse retinas. Likewise, Notch1 signaling pathway components such as Hes1 and Hes5, which also are essential for glial cell fate specification, were downregulated in CCN2-deficient retinas. Hes 1 expression was shown to be particularly important because Hes1 null mice can produce all six neuronal cell classes of the retina but lack MGCs (Zhu et al., 2013). Taken together, these observations indicate that CCN2 expression in mouse RPCs potentially determines their MGC fate, and possibly focuses RPC commitment to the glial cell lineage. Because CCN2-deficient mice die at birth and since the vast majority of MGCs are generated postnatally, further experiments will be needed to determine how CCN2 expression influences MGC formation and function postnatally and in adult mouse retina.

Regulation of specific gene expression is the endpoint of a network of signaling pathways enabling the cells to process signals they receive simultaneously from different receptors. The multitude of genes affected by CCN2 deletion suggests that CCN2 signals impacted global developmental pathways of retinogenesis. We found that Notch, Wnt, Shh, and Hippo YAP signaling pathways were specifically dysregulated in the retina upon CCN2 deletion in mice in line with the known functions of these pathways in nervous-system development (Dhillon and Dhillon, 2008). In retinal neuroblastic cells, CCN2 expression largely overlapped with that of YAP, the core component of the Hippo kinase signaling pathway. Downregulation of YAP expression was a direct consequence of CCN2 gene deletion in the retina suggesting that YAP is a downstream effector of CCN2 signals. Such regulatory relationship between CCN2 and YAP has been reported by Moon et al in the developing retinal vasculature as well (Moon et al., 2020). YAP plays a pivotal role in promoting cell cycle exit and terminal differentiation in various tissues including the CNS (Jin et al., 2020). As a transcriptional co-activator, YAP regulates the transcription of many genes involved in cell cycle entry and exit, tissue-specific cell differentiation and apoptosis (Hansen et al., 2015). Accordingly, YAP deletion resulted in decreased progenitor cell survival in the embryonic neuroepithelium, whereas its activation led to the opposite phenotype, namely increased progenitor and stem cell replication (Cao et al., 2008). In a different study, overexpression of YAP in the developing mouse retina enhanced RPC proliferation and inhibited their differentiation (Asaoka et al., 2014). YAP was shown to regulate the timing of PR differentiation through regulation of the expression of the Otx, Crx and rhodopsin genes (Asaoka et al., 2014). However, our data further established a bifunctional involvement of YAP in determining Sox9+ MGC differentiation as well. We found that Müller glia marked by Sox9 expression were nearly absent in retinal explants in culture, but their number increased by 70% upon ectopic expression of YAP or CCN2. Like CCN2, YAP expression persists in the adult retinal MGCs (Hamon et al., 2017) although its regulation and function in the adult eye is yet to be investigated. In the mouse retina, where MGCs do not spontaneously proliferate, the CCN2-YAP functional interaction potentially controls MGC quiescence state and response to injury. Further studies of the role of the CCN2-YAP regulatory axis in MGC homeostatic function and retinal degenerative diseases are warranted.

Moreover, YAP functions as a signaling nexus and integrator of several other signaling pathways critical for CNS development. These include the Notch, Wnt and Shh signaling pathways, all of which were altered in CCN2-deficient mice. Notch pathway components such as Dll1, Dll3, Hes 1, Hes 5, Hey1 and Hey 1 were particularly downregulated in the retina of CCN2-deficient mice. The expression of Hes1 and Hes5, which are inhibitors of neuronal differentiation, keep RPCs in an undifferentiated progenitor state or promote MGC differentiation during the late stage of retinal development (Zhu et al., 2013). Reduced levels of Hes1 and Hes5 corroborates our finding of a major function of CCN2 in gliogenesis. Furthermore, our data showed association between CCN2 and the canonical Wnt pathway. The Wnt/β-catenin pathway seemed to be activated in CCN2-deficient retinas through upregulation of Wnt2b and LRP5/6. Elevated levels of these Wnt components may have important consequences on RPC proliferation and competency. A study by Kubo et al (Kubo et al., 2005) showed that stable overexpression of Wnt2b in retinal explants inhibited cellular differentiation through downregulation of the expression of multiple proneural transcription factor genes. Although the effects of Wnt2b were reported to include promoting and prolonging proliferation of RPCs, such effects were not manifested in CCN2-deficient retina probably due to counterbalancing effects by other factors (e.g., Notch inhibition). In the canonical signaling pathway, Wnt proteins bind to Frizzled and LRP5/6, thereby activating Dishevelled, destabilizing Axin-1, and inactivating glycogen synthase kinase (GSK)-3β, which subsequently targets β-catenin to the cell nucleus to activate the transcription of target genes such as cyclin D1 (Liu et al., 2003). However, the functional significance of LRP5/6 upregulation during retinogenesis remains unclear. Recent studies suggested that the canonical Wnt pathway operates mostly in the marginal part of the optic vesicles where undifferentiated retinal progenitor cells reside and biases the RPC fate toward retinal pigment epithelium (RPE) (Fujimura, 2016). Future studies will investigate the effects of CCN2 deficiency on the growth and differentiation of other ocular tissue components (e.g., RPE).

Like Wnt, sonic hedgehog pathway components such as the transmembrane receptor patched (Ptch1), Gli1, and Sufu were upregulated upon CCN2-deletion in embryonic retina. Levels of Ptch and Gli1 proteins are commonly increased upon Shh activation (Rimkus et al., 2016). In rodents, recombinant Shh-N promotes retinal progenitor proliferation in cultures (Wall et al., 2009), and partial depletion of Shh decreases proliferation. Exposure of rat retinal cultured cells to the N-terminal recombinant Shh protein caused a transient increase in the number of RPCs, and an increase in the number of PRs (Wan et al., 2007). In the mouse retina, complete ablation of Shh signals profoundly influenced the fate determination of a subset of early-born neurons, primarily RGCs and cone PRs (Moshiri et al., 2005). However, whether CCN2 deletion induced full or partial activation of Shh signals remains to be determined. Further studies will be needed to assess the specific effects of CCN2 on Shh signaling pathway and its integration with other pathways during retinogenesis.

In summary, our molecular and genetic studies indicate that CCN2 signals affected both progenitor cell proliferation and lineage commitment. How might CCN2 coordinate these processes? Our data suggest a model in which CCN2 signals impact the cell cycle machinery in all progenitors, but ultimately critically influence the gliogenic fate, during which at least one postmitotic cells that maintains CCN2 expression is generated. Mature MGCs which persistently express CCN2 in the adult retina, exhibit the potential to dedifferentiate and become RPCs (Besser et al., 2012) suggesting a potentially important role of CCN2 in the RPC-MGC differentiation and dedifferentiation process. Future investigations will broaden our understanding of the role of CCN2 in the balanced production of diverse neuronal and non-neuronal cell types and maintenance of retinal tissue homeostasis during development and diseases.

## Materials and methods

### Transgenic and CCN2-deficient Mice

Animal studies were carried out in accordance with the recommendations in the Guide for the Care and Use of Laboratory Animals of the National Institutes of Health. Mice were handled and housed according to the approved Institutional Animal Care and Use Committee protocol 14-10425 of SUNY Downstate Medical Center. GENSAT Tg(*CTGF*-EGFP)FX156Gsat/Mmucd (011899-UCD) mice, referred to herein as CCN2-GFP mice carrying enhanced green fluorescent protein (GFP) under the control of the CCN2 promoter, were developed under the NINDS-funded GENSAT BAC transgenic project (Gong et al., 2003) and obtained from the Mutant Mouse Regional Resource Center. CCN2-GFP mice were initially in the FVB/N-Swiss Webster background and later backcrossed for >10 times in the C57BL/6J genetic background. CCN2^flox/flox^ were previously described (Liu et al., 2011). CMV-Cre mice which induces deletion of *loxP*-flanked genes in all tissues including germ cells (Schwenk et al., 1995) were from Jackson Laboratory. Mice with germline deletion of CCN2 were generated by cross-breading CCN2^flox/flox^ with CMV-Cre mice to produce CCN2^flox/+^ CMV-Cre^-/-^ and CCN2^flox/+^ CMV-Cre^+/-^. The latter were further crossed among each other or with CCN2^flox/flox^ to produce CCN2^flox/+^CMV-Cre^+/-^, CCN2^flox/+^CMV-Cre^-/-^ or CCN2^flox/flox^CMV-Cre^+/-^mice (i.e, CCN2^-/-^). Genotyping was determined by qPCR to identify mice with floxed alleles, and Cre allele (CCN2^-/-^) and homozygous floxed alleles (CCN2^-/-^). Recombination levels in CCN2^-/-^ mice as compared to WT were determined as described in the Results section. Retinas from mice with the CMV-Cre, CCN2^flox/+^ or CCN2^flox/flox^ genotype were undistinguishable in the assays described herein and were used as controls (i.e., CCN2^+/+^) for CCN2^-/-^ retinas. Since gender has not previously been reported as a biological variable for retinal cell differentiation, embryos and mice of both genders were used undistinguishably in our studies. Ages of embryos and mice used in experiments are as indicated in the figures. A minimum of three embryos in each group was analyzed.

### Retinal explant culture and lentivirus transduction

Eyecups containing the neuroretina and lens were dissected from E14 CCN2-GFP, CCN2^+/+^ and CCN2^-/-^ mouse embryos, and placed into a 96-well plate in 100 μl of MEM-HEPES supplemented with 10% heat-inactivated fetal bovine serum, 200 μM L-glutamine, 5.75-mg/ml glucose, 100-U/ml penicillin, and 100-μg/ml streptomycin (Invitrogen). The plate was maintained in Hera Cell culture incubator (37 °C, 5% CO2) for 24 hrs. Thereafter, a lentiviral vector carrying the GFP gene or CCN2 or YAP cDNA (Addgene) expressed from the cytomegalovirus (CMV) promoter was added to the culture medium. Lentiviruses were produced by transient CaPO4-mediated transfection of 293T cells with lentiviral backbone plasmids, a vsv.g-envelope plasmid, and a gag/pol plasmid. Titers of 5 × 10^6^ IU/ml were achieved and concentrated by ultracentrifugation to 1 × 10^12^ IU/ml. Fresh new medium was added every 48 h. After 6 days, the eyecups were fixed in 4% paraformaldehyde (PFA) in PBS for 2 hr at room temperature, and retinas were dissected and processed for cryosections. Three eyecups in each group were analyzed.

### Immunohistochemistry

Embryos were collected from timed pregnant CCN2^flox/flox^ CMV-Cre^+/-^ mice and genotyped as previously described (Moon et al., 2020). Eye was enucleated and placed in 4% PFA for 30 min. after a s series of washes with PBS and incubation in 10-30% sucrose solutions, eyes were frozen in Tissue-Tek Optimal Cutting Temperature (OCT) compound and stored at -80 °C until further use. In separate sets of eyes, retinas were dissected and then frozen in OCT. Transverse frozen sections (10 mm) were used for immunohistochemistry using standard protocols. Tissue processing and immunostaining were carried out as described previously (Lee et al., 2019). Tissue sections were then permeabilized in 0.1% Triton X-100 at room temperature for 20 min and further single or double stained with primary and secondary antibodies. The following primary antibodies were used: anti-Brn3a/Pou4f1 (mouse, Millipore, MAB1585, 1:500); anti-Chx10 (goat, Santa Cruz, sc-21690, 1:2000); anti-Pax6 (rabbit, Biolegend, 901301, 1:1000); anti-Sox9 (rabbit, Millipore, AB5535, 1:800); anti-Sox2 (rabbit, Millipore, AB5603, 1:500), rabbit anti-recoverin (rabbit, Millipore, AB5585, 1:200), anti-Islet 1&2 (mouse, DSHB, 1:50), anti-Ki67 (rabbit, Abcam, ab16667, 1:100), and isolectin B4 (5 mg/ml, Vector). The secondary antibodies are as follows: Alexa fluor donkey anti-mouse 488 (Invitrogen, A21202, 1:500); Alexa fluor donkey anti-mouse 594 (Invitrogen, A21203, 1:500); and Alexa fluor donkey anti-rabbit 594 (Invitrogen, A21207, 1:500). Sections were washed three times (5 mins each) with 1X PBS at room temperature, and then mounted with 4′,6-Diamidino-2-phenylindole dihydrochloride (DAPI) for nuclear counterstaining. Images were captured using a Leica DM5000 B fluorescence imaging system (Leica).

### Fluorescence-activated cell sorting (FACS)

Retinas from E14 CCN2-GFP mice were dissociated and live cells were stained for 30 min at 37°C with Vibrant Violet dye (Invitrogen) and then subjected to FAC analysis using a CyAn flowcytometer (DAKO Cytomation). DNA content was determined by gating on EGFP-positive cells and assessing Vibrant Violet fluorescence. DNA Content histograms were modeled for cell cycle parameters in 30,000 cells using MODFIT software.

### 5-Bromo-2′-deoxyuridine/BrdU incorporation assay

Newborn cells in the retina were labeled through intraperitoneal injection of BrdU (10 mg/kg of body weight) and collected labeled retinas were treated with 2M HCl for 30 mins at 37° C for antigen retrieval. Retinas were further incubated for 10 min with 0.1 M sodium borate buffer (pH 8.5) at room temperature. BrdU was detected using directly conjugated mouse anti-BrdU Alexa 488 (Molecular Probes, A21202; 1:500 dilution). The number of BrdU-positive cells in equivalent areas of retinal sections was determined.

### Terminal deoxynucleotidyltransferase-mediated dUTP-biotin nick end labeling (TUNEL) staining

TUNEL staining was performed with an ApopTag fluorescein *in situ* apoptosis detection kit (EMD Millipore) according to the manufacturer’s protocol. TUNEL-positive nuclei were counted, and the number was normalized to the number of DAPI-positive nuclei.

### RNA isolation and RT-qPCR

Total RNA was extracted from mouse retina using TRIzol reagent (Sigma). The cDNA synthesis was carried out by using Prime script reverse transcript kit (Takara). Highly specific primers were designed using Web-based primer design programs. A qPCR was performed with biological triplicates using Power SYBR™ green PCR premix (Applied biosystems). The cycling parameters for qPCR amplification reactions were: AmpliTaq activation at 95°C for 10min, denaturation at 95°C for 15s, and annealing/extension at 60°C for 1min (40 cycles). Triplicate *Ct* values were analyzed with Microsoft Excel using the comparative *Ct* (ΔΔ^Ct^) method as described by the manufacturer. The transcript amount (determined by the −2^ΔΔCt^ threshold cycle [CT] method) was obtained by normalizing to an endogenous reference (18S rRNA) relative to a calibrator.

### Transcriptome profiling using RNA-seq

Retinas were harvested (at ∼10:00am) from E18.5 CCN2^+/+^ and CCN2^-/-^ from 3 different pregnant dams (3 replicates from each control and mutant mice were used). One pair of retinas from each embryo were rapidly dissected out of the eye cup on ice and stored in Trizol at -80°C. RNA from one pair of retinas was individually isolated, with three pairs of retinas in total for WT and mutant mice. The retinas were homogenized with an electric pellet pestle motor in 500uL of Trizol. The quantity and quality of RNA were verified with NanoDrop 1000 (Thermo Scientific, DE). RNA-Seq analysis was performed at GENEWIZ following the manufacturer’s protocol. Strand-specific mRNA-seq libraries for the Illumina® NovaSeq™ platform were generated using high-quality total RNA as input for the dUTP library preparation method. Briefly, the mRNA fraction was purified from total RNA by polyA capture, fragmented and subjected to first-strand cDNA synthesis with random hexamers in the presence of Actinomycin D. The second-strand synthesis wasperformed incorporating dUTP instead of dTTP. Barcoded DNA adapters were ligated to both ends of the double-stranded cDNA and subjected to PCR amplification. The resultant library was checked on a Bioanalyzer (Agilent) and quantified. The libraries were multiplexed, clustered, and sequenced on the Illumina HiSeq platform. The sequencing run was analyzed with the Illumina CASAVA pipeline (v1.8.2), with demultiplexing based on sample-specific barcodes. 20-30 million read pairs per sample were perfoemd. The raw sequencing data produced was processed removing the sequence reads which were of too low quality (only ‘‘passing filter’’ reads were selected) and discarding reads containing adaptor sequences or PhiX control with an in-house filtering protocol.

### Filtering of reads and pathway analysis

Image analysis and base calling were performed using Solexa pipeline v1.8 (Off-Line Base Caller software, v1.8). Sequence quality was examined using the FastQC software. The trimmed reads (trimmed 5’,3’-adaptor bases using Cutadapt) were aligned to reference genome using Hisat2 software (v2.0.4) (Kim et al., 2015). The transcript abundance for each sample was estimated with StringTie (v1.2.3) (Pertea et al., 2015), and the Fragments Per Kilobase of transcript per Million mapped (FPKM) value (Mortazavi et al., 2008) for gene and transcript levels were calculated with R package Ballgown (v2.6.0) (Frazee et al., 2015). The thresholds used for identifying differentially expressed genes (DEGS) were: (i) DESeq. 2 mean normalized counts >10; (ii) padj-value <0.5, and (iii) log2fold change >0 (Love et al., 2014). rMATS (Shen et al., 2014) was used to analyze alternative splicing events.The differentially expressed genes/transcripts were analyzed for their enrichment in gene ontological functions or pathways using the TopGo R software package. The statistical significance of enrichment was given as p-value by Fisher exact test and -log10(p) transformed to Enrichment score. Principal Component Analysis (PCA) was performed with genes that have the ANOVA p value *<* 0.05 on FPKM abundance estimations. Statistical or graphics computing, correlation analysis, Hierarchical Clustering, scatter plots and volcano plots were performed in R, Python or shell environment. The raw RNA-Seq data and analyzed FPKM values can be accessed through NCBI/GEO (GSE146900).

### Gene set enrichment analysis (GSEA)

RNA-Seq dataset expression of gene profiling files downloaded from the dataset was analyzed by Gene Set Enrichment Analysis (GSEA, http://www.broad.mit.edu/gsea/index.html). The whole genome (27455 genes) with expression values were uploaded to the software and compared with catalog C5 gene ontology gene sets in MsigDB (Subramanian et al., 2005), which contains 233 GO cellular component gene sets, 825 GO biological process gene sets, and 396 GO molecular function gene sets. GSEA was run according to default parameters collapses each probe set into a single gene vector (identified by its HUGO gene symbol), permutation number = 1000, and permutation type = “gene-sets”. Calculation of the false discovery rate (FDR) was used to correct for multiple comparisons and gene set sizes.

### Western immunoblotting

For retinal protein analysis, mouse eyes were enucleated, and the retinas were carefully dissected and homogenized in lysis buffer containing 1% Triton X-100, a 1% volume of phosphatase, and a protease inhibitor mixture. Protein samples (25 μg) were fractionated in a 10% SDS-polyacrylamide gel and transferred to a nitrocellulose membrane, and Western blot analysis was performed by incubation with primary CCN2 antibodies (Moon et al., 2020) and then secondary antibodies. Immunodetection was performed using enhanced chemiluminescence (Pierce).

### Quantitative and Statistical Analyses

For CCN2-deficient mice, at least three retinas from littermates were analyzed for each type of control and mutant animals. Counting of specific marker-labeled cells was performed on 3-6 random non-overlapping fields in the intermediate region for each retina. The number of marker labeled cells was normalized to the total number of DAPI-labeled nuclei. For GFP-positive or overexpression experiments, a few hundred maker-positive cells were counted in at least 3 retinas. Data were expressed as means ± SE. To test differences among several means for significance, a one-way analysis of variance with the Newman-Keuls multiple comparison test was used. Where appropriate, a *post hoc* unpaired *t* test was used to compare two means/groups. Non-parametric Mann-Whitney or Kruskal-Wallis test was used when data collected do not have normal or clear distribution. Statistical significance was set to *p* value less than 0.05.

## Acknowledgments

We are thankful to Dr. Stephen J. Weiss (University of Michigan) for his involvement in the project conception. We appreciate the technical contribution of Sohyun Moon and Xin Chen. We thank all past and present lab members for their contributions to the generation and characterization of genetically modified and transgenic animals and discussions during the preparation of the manuscript. This work was supported in part by grants from the National Eye Institute of the National Institutes of Health (grant EY024998) to B.C.

## Author contributions

G.M. performed in vivo deletion of CCN2 and retinal phenotype analyses; G.L. and A.B. performed mouse genotyping, slide microscope scanning and blinded analyses; J.S. performed RNA seq and GSEA analyses; M.E.H. contributed to data interpretation and discussion; B.C. designed the entire project and wrote the paper.

## Data Availability

RNA-seq data have been deposited at the NCBI Gene Expression Omnibus under the series record GSE171232.

